# Does the punishment fit the crime? Consequences and diagnosis of misspecified detection functions in Bayesian spatial capture-recapture modelling

**DOI:** 10.1101/2021.01.12.426298

**Authors:** Soumen Dey, Richard Bischof, Pierre P. A. Dupont, Cyril Milleret

## Abstract

- Spatial capture-recapture (SCR) is now used widely to estimate wildlife densities. At the core of SCR models lies the detection function, linking individual detection probability to the distance from its latent activity center. The most common function (half-normal) assumes a bivariate normal space use and consequently detection pattern. This is likely an oversimplification and misrepresentation of real-life animal space use patterns, but studies have reported that density estimates are relatively robust to misspecified detection functions. However, information about consequences of such misspecification on space use parameters (e.g. home range area), as well as diagnostic tools to reveal it are lacking.
- We simulated SCR data under six different detection functions, including the half-normal, to represent a wide range of space use patterns. We then fit three different SCR models, with the three simplest detection functions (half-normal, exponential and half-normal plateau) to each simulated data set. We evaluated the consequences of misspecification in terms of bias, precision and coverage probability of density and home range area estimates. We also calculated Bayesian p-values with respect to different discrepancy metrics to assess whether these can help identify misspecifications of the detection function.
- We corroborate previous findings that density estimates are robust to misspecifications of the detection function. However, estimates of home range area are prone to bias when the detection function is misspecified. When fitted with the half-normal model, average relative bias of 95% kernel home range area estimates ranged between −25% and 26% depending on the misspecification. In contrast, the half-normal plateau model (an extension of the half-normal) returned average relative bias that ranged between −26% and −4%. Additionally, we found useful heuristic patterns in Bayesian *p*-values to diagnose the misspecification in detection function.
- Our analytical framework and diagnostic tools may help users select a detection function when analyzing empirical data, especially when space use parameters (such as home range area) are of interest. We urge development of additional custom goodness of fit diagnostics for Bayesian SCR models to help practitioners identify a wider range of model misspecifications.

## 1 Introduction

Spatial capture recapture (SCR) models are used to estimate the density of wildlife populations (see Royle et al., 2014; Borchers and Fewster, 2016) and, increasingly, other important ecological parameters (Efford et al., 2016). These are hierarchical models that use the spatial information from repeated individual encounters to estimate the location of individual activity centers, while accounting for imperfect detection. The Bayesian paradigm and accessible programming languages (de Valpine et al., 2017; Plummer, 2003) provide a convenient framework for developing and fitting customized SCR models (Bischof et al., 2020b; Efford and Schofield, 2020; Augustine et al., 2019; Turek et al., 2020), and we are experiencing a surge of innovations in Bayesian SCR models of growing scope and complexity (Royle et al., 2018). Conspicuously lagging behind, are studies that systematically evaluate the consequences and discuss the diagnostics of common model misspecifications (Dupont et al., 2019).

In this study, we focus on the consequences and diagnosis of misspecifications of a core component shared by all SCR models: the detection function. Detection probability in SCR analysis is modelled as a function of the distance between the latent locations of individual activity centers (ACs) and the locations of the detectors. The shape of the detection function can be interpreted as a reflection of an individual’s space use around its activity center. The half-normal detection function (HN) is the most common detection function in SCR modelling and assumes a bivariate normal model for animal space use, therefore characterising a monotonic decay in detection probability with increasing distance from the AC (Efford, 2004). However, animal space use and home range configurations can vary substantially between (Gittleman and Harvey, 1982; Ofstad et al., 2016) and even within species (Kie et al., 2002; Efford and Mowat, 2014). It is therefore reasonable to assume that ‘one size does not fit all’, as far as the detection function in SCR models is concerned. For example, territorial species may spend a disproportionate amount of time patrolling and scent marking the boundaries of their territory (Langergraber et al., 2017; Potts et al., 2013), leading to a bimodal or donut-shaped space use profile. Conversely, species or demographic groups with long exploratory forays (Zeale et al., 2012), may exhibit long-tailed space use distributions. Species with relatively even space use throughout a clearly defined home range (Pearce et al., 2013) may represent another deviation from the half-normal model, exhibiting a plateau in the utilization distribution followed by more or less rapid declines in utilization near the edge of the home range.

There are indications that density estimates produced by SCR models are reasonably robust to misspecifications of the detection function (Efford, 2004; Efford and Dawson, 2009; Russell et al., 2012). However, misspecifications may have consequences for estimates of other important parameters besides density/population size, such as home range area, habitat use/selection, movement, and connectivity, all of which feature increasingly in the SCR literature (Efford et al., 2016; Royle et al., 2013b; Gardner et al., 2018; Royle et al., 2013a). Furthermore, even in the absence of systematic bias, misspecifications could impact the confidence one can place in those estimates by affecting the associated precision and coverage probability (Bischof et al., 2020a).

While simulations can inform about the potential consequences of model misspecification, they cannot be used to identify misspecifications in empirical situations. Goodness of fit testing offers a formal tool for diagnosing violations of assumptions and is an important part of statistical analysis as it reduces the risk of drawing erroneous inference (Pradel et al., 2005). Bayesian *p*-values are frequently used to assess the goodness of fit in Bayesian modelling and do so by measuring the systematic dissimilarity between observed data and the posterior distribution (Gelman et al., 2014). Although Bayesian *p*-values have been used previously in SCR studies for assessing goodness of fit of different components of the model (Russell et al., 2012; Ergon and Gardner, 2014; Proffitt et al., 2015), their efficacy in diagnosing misspecified detection functions in SCR models has yet to be explored.

Using simulations, we evaluated the consequences of detection function misspecifications on key SCR parameter estimates and assessed whether Bayesian *p*-values can be used as a diagnostic tool to detect such misspecifications. First, we quantified the impact of the choice of detection function - relative to the function describing true individual space use - on SCR-derived estimates of density and home range area. Then, we calculated and compared a suit of Bayesian *p*-values and assessed their ability to reveal misspecifications. We discuss the implications of our results in the context of the choices and challenges faced by practitioners using SCR to analyze empirical data.

## 2 Methods

### 2.1 General approach

We conducted a simulation study to assess the consequences of misspecifying the detection function in SCR models. Six different detection functions were used to simulate contrasting animal space use patterns (in terms of profile shape and complexity) and generate corresponding spatial capture-recapture data (Fig. 1). We then fitted three SCR models differing in their detection function (half-normal, exponential or half-normal plateau; see Section 2.3.2 below) to the simulated data sets. We assessed the robustness of the models by calculating the relative bias, coefficient of variation, and coverage probability of the 95% credible intervals for the population size and home range area estimates (50% and 95% of the space use distribution). Finally, we calculated a series of Bayesian *p*-values and compared their ability to diagnose misspecifications of the detection function.

**Figure 1:**
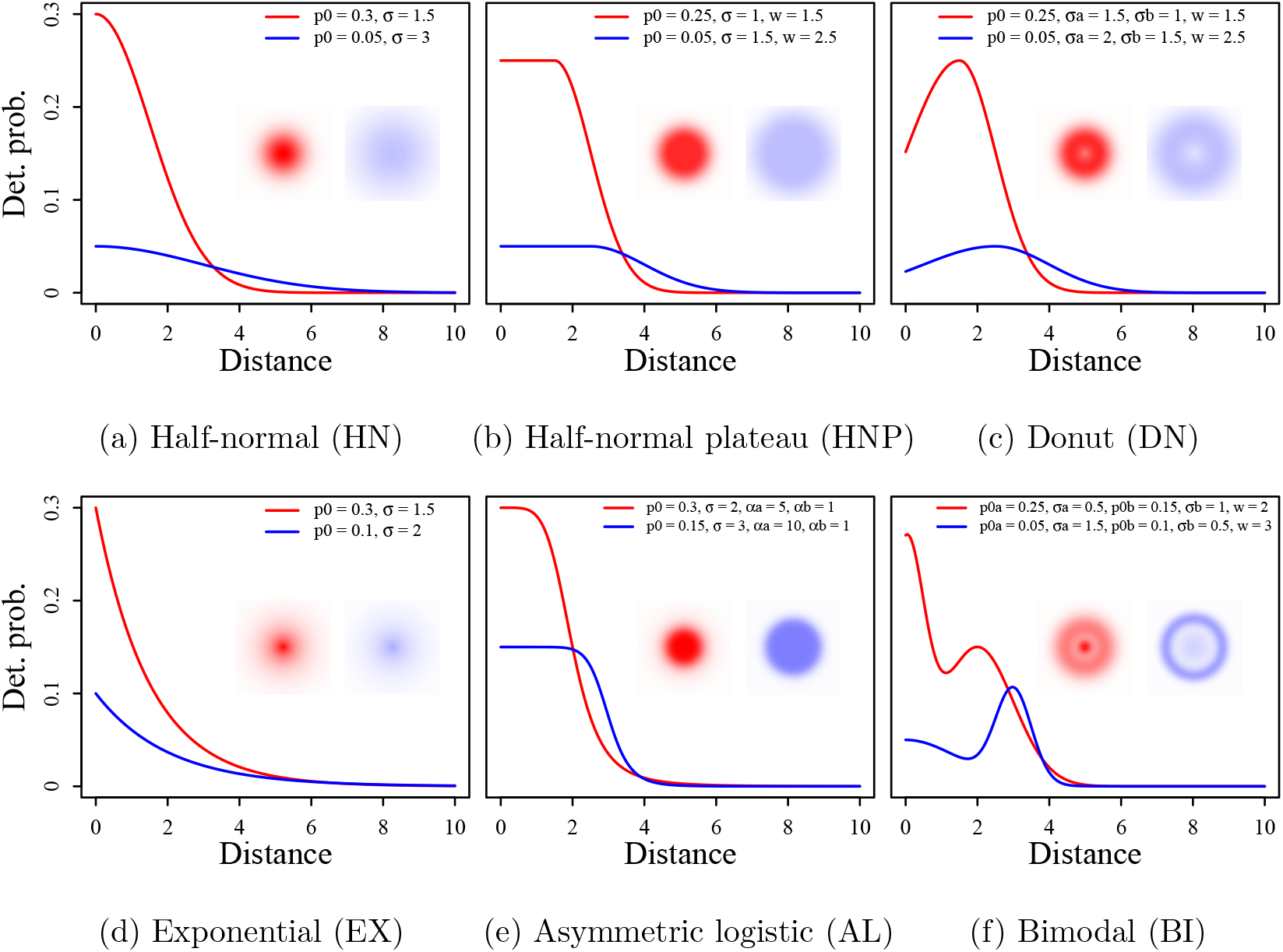
Visualization of six different detection functions (detection probability as a function of distance between the detector and individual activity center), both as kernel density profiles and raster maps. Realization of two different parameter sets are shown for each detection function, with red lines and shading correspond to parameter set 1, whereas blue lines and shading correspond to parameter set 2. Values of applicable detection function parameters (see main text for descriptions) are provided in the legend of each plot. Distances are provided in arbitrary distance units (du).

### 2.2 SCR model description

A single-season SCR model forms the basis for data simulation and model fitting in this study. SCR models are hierarchical in their structure with a component describing the spatial distribution of individuals in a given area (habitat) and an observation model describing individual detection probability in space, conditional on their AC location.

Consider a population of *N* individuals that reside in a bounded habitat 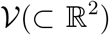 where each individual is assumed to move randomly around its AC, viz., 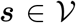. These latent (unobserved) ACs are assumed to be a realization of a homogeneous Poisson point process.

We used the data augmentation approach to model the number *N* of individuals in the habitat 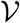 (Royle et al., 2009). We implemented this approach by choosing a large integer *M* to bound *N* and introduced a vector of *M* latent binary variables ***z*** = (*z*_1_*, z*_2_*, …, z_M_*) in the SCR model. We define *z_i_* = 1 if individual *i* is a member of the population and *z_i_* = 0 otherwise. We then assume that each *z_i_* is a realisation of a Bernoulli trial with parameter *ψ*, the inclusion probability.

An array of *J* detectors is located within the habitat 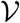. These detectors are kept active for one sampling occasion. The recorded observations (*y_ij_*’s) are taken as binary outcome of the capture-recapture survey. In other words, *y_ij_* = 1 if the *i*-th individual is detected at the *j*-th detector and *y_ij_* = 0 otherwise. We assume that *n* individuals are detected during the capture-recapture survey. Consequently, ***Y*** _obs_ is of dimension *n × J*. As part of the data augmentation approach, the SCR data set ***Y*** _obs_ = ((*y_ij_*)) is supplemented with a large number of “all-zero” encounter histories. The zero-augmented data set ***Y*** (dimension *M × J*) is constructed as follows:

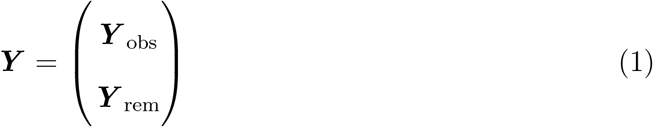

where ***Y*** _rem_ denotes the array of “all-zero” encounter histories with dimensions (*M − n*) *× J*.

A Bernoulli model, conditional on *z_i_*, is assumed for each observation *y_ij_*:

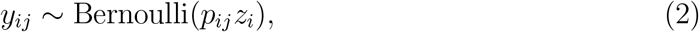

where *p_ij_* denotes the detection probability of the *i*-th individual at the *j*-th detector. The detection probability *p_ij_* is modelled as a function of the euclidean distance *d_ij_* between the detector location ***x** _j_* and individual AC location ***s**_i_*

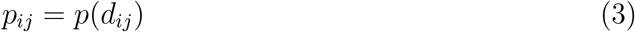

where

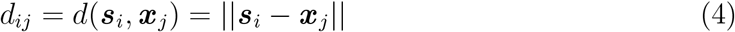

This formulation of *p_ij_* allows incorporation of individual and detector-level heterogeneity in detection probability. In addition, this implementation allows the user to specify the shape of the detection probability function *p* in terms of *d_ij_*.

### 2.3 Simulation design

#### 2.3.1 Habitat and detectors

We assumed a square habitat 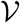 of 29 × 29 distance units (du). Centered on the habitat, is a 20 × 20 detector array (*J* = 400 detectors) with 1 du spacing between neighboring detectors. This configuration results in a 5-du un-sampled habitat buffer around the outermost detectors (Appendix Fig. 1)

#### 2.3.2 Detection functions

As noted in Section 1, we assume that the detection probability function reflects assumptions about individual home range space use around the AC. Space use varies between species, populations, and even individuals and there are numerous detection functions corresponding to the numerous space use distributions. For the purpose of this study, we limited ourselves to six functions, that differed substantially in the shapes they can potentially assume (Table 1 and Fig. 1). In SCR modelling, it is standard practice to assume that individual space use is circular and that it is highest at the AC location and gradually decays with distance from AC (e.g. Dorazio, 2013). Both the *half-normal* (HN) and *exponential* (EX) detection functions (Table 1 and Figs 1(a) and 1(d)) reflect this behaviour and are frequently used in SCR studies, with the half-normal being the most popular choice. In these two functions, *p*_0_ denotes the baseline detection probability (i.e., when the detector is notionally placed at the AC location; euclidean distance *d* = 0) and the scale parameter *σ* quantifies the spatial extent of animal space use around its AC.

**Table 1:**
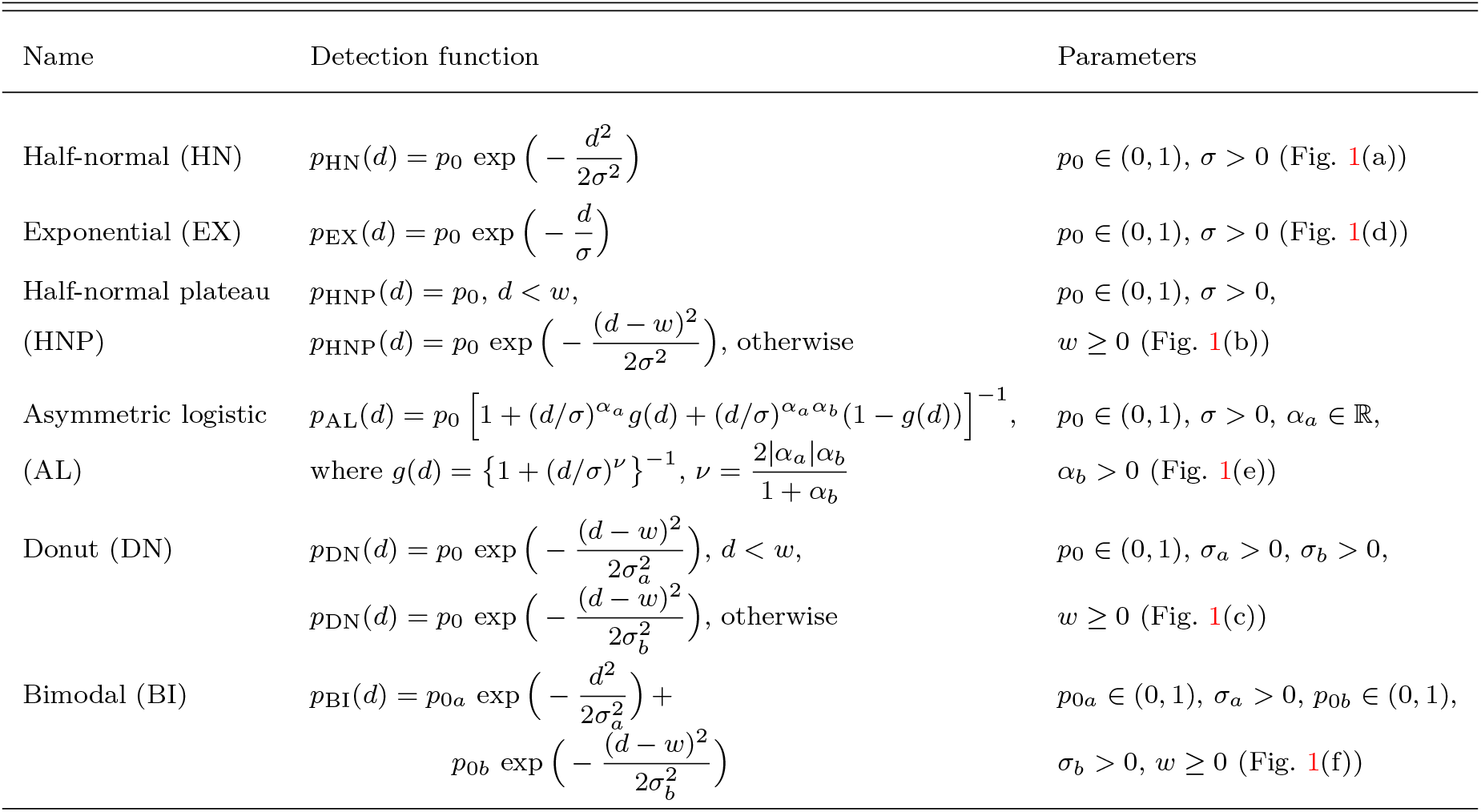
Detection probability functions

To allow greater flexibility in individual space use, we introduce two detection functions that permit uniform space use up to a certain radius and decays afterwards: the *Half-normal plateau* (HNP) and *Asymmetric logistic* (AL) (Table 1 and Figs 1(b) and 1(e)). The half-normal plateau is an extension of the half-normal detection function with an additional non-negative parameter *w* for the radius of uniform activity. By contrast, the asymmetric logistic detection function is an extension of the logistic curve model with an additional parameter *α_b_* for the second curvature, which allows the detection function to be asymmetrical and allows for added plasticity in HR space utilization (Ricketts and Head, 1999). The other parameters of this function are *p*_0_, the baseline detection probability, *α_a_*, the first curvature parameter and *σ*, the distance from the AC where the asymmetric logistic detection function takes the value *p*_0_*/*2 (Richards, 1959).

We also used two detection functions that display an increase in space use with distance from AC (e.g., common pipistrelle bats: *Pipistrellus pipistrellus*, Nicholls and Racey, 2006): the *donut* (DN) and *bimodal* (BI) detection functions (Table 1 and Fig. 1(c) and (f)). The donut function has a single peak with highest detection probability *p*_0_ at a distance *w* from AC. The two tails on the two sides of the peak correspond to two bivariate normal density curves with the same mean *w* but with different scale parameters *σ_a_* and *σ_b_*. The bimodal function is a stochastic mixture of two univariate normal densities with means 0 and *w*(*>* 0) and scale parameters *σ_a_* and *σ_b_*, with mixture weights *p*_0*a*_ and *p*_0*b*_, respectively.

### 2.4 Scenarios

Spatial capture recapture data sets were simulated assuming a single season SCR model (Section 2.2) for the six detection functions using two sets of parameters for each (Fig. 1 and Table 2). This resulted in 12 different simulation scenarios where the total number of individuals with ACs in the habitat were kept constant at *N* = 200. The two parameter sets were chosen to result in a similar number of detected individuals (around 120-140) but different HR area and different total detection counts (i.e., 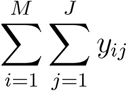; Appendix Fig. 2). Due to computation limitations, we could not fit SCR models with all six detection functions used during simulation (see Section 2.3.2). Instead we fitted the two most common ones: half-normal (HN) and exponential (EX) as well as the half-normal plateau (HNP) for its expected flexibility. We fitted each of the three models to separate 50 random realizations (repetitions) of each of the 12 scenarios, resulting in 1800 simulated data sets.

**Table 2:** Parameter values of the six detection functions used for simulating spatial capture-recapture data. Also shown are the corresponding 95% quantile home range area, no. of detected individuals (mean, 2.5% and 97.5% quantiles) for two parameter sets associated with smaller (set 1) and larger (set 2) home range sizes.

| Detection functions | Parameter set 1 |  |  | Parameter set 2 |  |  |
| --- | --- | --- | --- | --- | --- | --- |
|  | Parameter values | 95% HR area (du sq.) | No. of detected individuals | Parameter values | 95% HR area (du sq.) | No. of detected individuals |
| Half-normal (HN) | $p_0 = 0.3, \sigma = 1.5$ | 42.35 | 123 (111, 137) | $p_0 = 0.05, \sigma = 3$ | 169.40 | 123 (110, 137) |
| Exponential (EX) | $p_0 = 0.3, \sigma = 1.5$ | 158.87 | 136 (123, 149) | $p_0 = 0.1, \sigma = 2$ | 274.48 | 117 (104, 132) |
| Half-normal plateau (HNP) | $p_0 = 0.25, \sigma = 1$<br>$w = 1.5$ | 39.50 | 133 (123, 146) | $p_0 = 0.05, \sigma = 1.5$<br>$w = 2.5$ | 95.75 | 126 (114, 138) |
| Asymmetric logistic (AL) | $p_0 = 0.3, \sigma = 2$<br>$\alpha_a = 5, \alpha_b = 1$ | 57.82 | 127 (113, 141) | $p_0 = 0.15, \sigma = 3$<br>$\alpha_a = 10, \alpha_b = 1$ | 40.92 | 127 (115, 139) |
| Donut (DN) | $p_0 = 0.25, \sigma_a = 1.5$<br>$\sigma_b = 1, w = 1.5$ | 39.98 | 133 (120, 146) | $p_0 = 0.05, \sigma_a = 2$<br>$\sigma_b = 1.5, w = 2.5$ | 97.14 | 125 (115, 137) |
| Bimodal (BI) | $p_{0a} = 0.25, \sigma_a = 0.5$<br>$p_{0b} = 0.15, \sigma_b = 1,$<br>$w = 2$ | 49.64 | 133 (120, 147) | $p_{0a} = 0.05, \sigma_a = 1.5$<br>$p_{0b} = 0.1, \sigma_b = 0.5,$<br>$w = 3$ | 46.81 | 121 (107, 136) |

### 2.5 Model fitting

We fitted all models using Markov chain Monte Carlo (MCMC) simulations with NIMBLE (de Valpine et al., 2017) in R (R Core Team, 2019). To reduce computation time, we implemented the local evaluation approach (Milleret et al., 2019). We ran three chains of 15 000 iterations including an initial burn-in phase of 5000 iterations. MCMC convergence of each model was monitored using the Gelman-Rubin convergence diagnostics 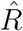 (also known as potential scale reduction factor, Gelman et al., 2014).

During preliminary analyses, we observed slow mixing of the Markov chains for parameters *σ* and *w* of the HNP detection function with the standard MCMC within the Gibbs sampler. To improve mixing, we used the recycling Gibbs sampler (Martino et al., 2018) for these parameters. R code for implementing simulations and fitting the single-season SCR model with different detection functions is provided in the supplementary material.

### 2.6 Consequences of misspecification

#### 2.6.1 Deriving population size *N*

In most Bayesian SCR models, population size is a derived parameter which follows a binomial distribution with parameters *M* and *ψ*, 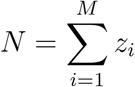. The parameter *ψ* gives the probability that an arbitrary individual from of the set of *M* individuals is a member of the population. For model fitting, we set *M* at 400 for all scenarios. We assigned uniform priors to all probability parameters (e.g., *p*_0_, *ψ*) with the support (0, 1). In addition, we assumed strictly positive uniform priors for the other model parameters (e.g., *σ*, *w*).

#### 2.6.2 Deriving home range area

Because the parameters from the different detection functions have different meanings and can not be directly compared, we based our comparison of the different models on the estimates of home range areas that can be derived from the detection function parameter estimates. Each of the detection functions *p*(***x**, **s***) used during fitting (Table 1) is proportional to the probability density function of a certain bivariate distribution *g*(***x** | **s***) and individual detection locations can be thought of as draws from this bivariate distribution

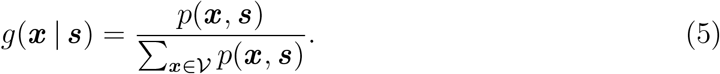

For any such space use probability distribution, we can find the quantile *r_α_* such that *α*% of all movements lie within the circle of radius *r_α_* centered on ***s***. We can also denote *A_α_*, the *α*% home range area, as the set of all points 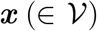 such that *||**x** − **s**|| ≤ r_α_*. Assuming a circular home range area, *A_α_* is then simply calculated as 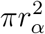. The home range area can therefore be derived by finding *r_α_* such that 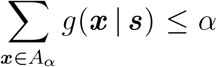. Here, we used a bisection algorithm to find the root of the above optimization problem (i.e., *r_α_*; see supplementary material for the R code to derive home range area) (Corliss, 1977).

For the half-normal detection function *p*_HN_(***x**, **s***) = *p*_0_ exp(*−*0.5*σ^−^*^2^*||**x** − **s**||*^2^), an analytical solution exists to calculate *r_α_*. Since *σ^−^*^2^*||**x** − **s**||*^2^ follows a chi-square distribution with 2 degrees of freedom, *r_α_* can be calculated as 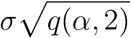 where *q*(*α,* 2) is the *α*% quantile of a chi-square distribution with 2 df (see Royle et al., 2014). As the analytical solution is more accurate and faster to obtain than numerical approximations, we analytically derived home range radius and area for the HN model but used numerical approximations for the other five detection functions for which no simple analytical solution exists. In order to compare the different models, we calculated home range radius and area for two *α* levels (50% and 95%) using a thinned sample (thinning rate = 10) combining all the MCMC chains, thus producing posterior distributions of the 50% and 95% home range radius and area (Table 2).

#### 2.6.3 Model performance measures

We used relative bias, coefficient of variation and coverage probability to evaluate the effect of detection function misspecifications on the estimates of population size and home range area. Suppose {*θ*^(*r*)^: *r* = 1, 2*, …, R*} denotes a set of MCMC draws from the posterior distribution of a scalar parameter *θ*.

##### Relative bias

Relative bias (RB) is calculated as

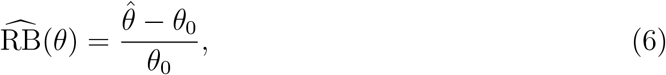

where *θ*_0_ is the true (simulated) value of a scalar parameter *θ* and 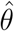 denotes the posterior mean estimate which is computed as 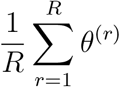.

##### Coefficient of variation

The precision of a parameter *θ* was measured by its coefficient of variation (CV) estimate:

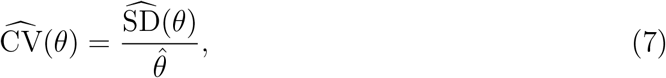

where 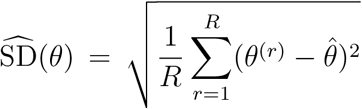 is the posterior standard deviation estimate of the parameter *θ*.

##### Coverage probability

Coverage probability was computed as the proportion of model fits (to the 50 simulations within a set, see Section 2.4) for which the estimated 95% credible intervals (CI) of *θ* contains the true value *θ*_0_.

### 2.7 Goodness of fit testing

We assessed the goodness of fit of each SCR model fitted to a simulated data set using the Bayesian *p*-value approach (Gelman, 2003). Bayesian *p*-values are calculated by using a test quantity *T* to measure the discrepancy between the observed data ***Y*** and their posterior replicates ***Y*** ^rep^. In practice, the posterior replicates of a SCR data set ***Y*** are obtained by drawing one 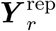 from its assumed model for each posterior simulation of parameter vector ***μ**_r_*, *r* = 1, 2*, …, R*. Consequently, two samples (*T* (***Y**, **μ***_1_)*, T* (***Y**, **μ***_2_)*, …, T* (***Y**, **μ**_R_*))’ and 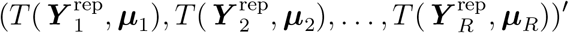 are generated. Then the Bayesian *p*-value is calculated as the proportion of times the replicated test quantity 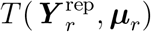 is greater than the observed quantity *T* (***Y**, **μ**_r_*):

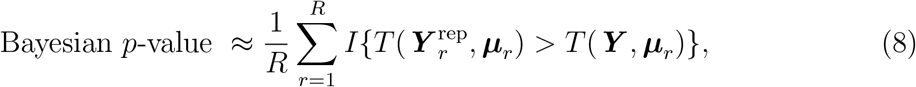

where *R* denotes the number of MCMC samples and *I*(*x*) is an indicator function taking the value 1 if *x* is true and 0 otherwise. For a good fit, the Bayesian p-value should be near 0.5 rather than the extremes 0 or 1.

Currently, a universal test quantity *T* is not available for SCR models. We implemented four different test quantities *T* to compare their ability to identify misspecifications in the detection function. We used the Freeman-Tukey (FT) measure (Freeman and Tukey, 1950) for its applicability for sparse data sets due to the variance stabilizing square root transformation (Brooks et al., 2000):

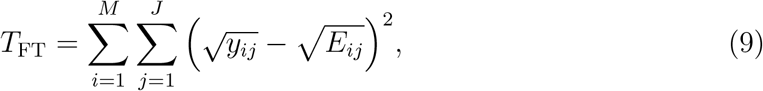

where *y_ij_* and *E_ij_* denote the capture-recapture observation and its expected value under the model for individual *i* at the *j*-th detector location, respectively. We also used two versions of this statistic pooled at the individual or detector level:

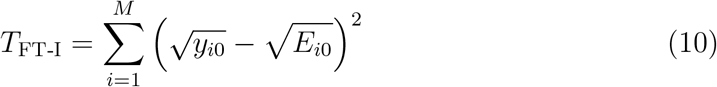

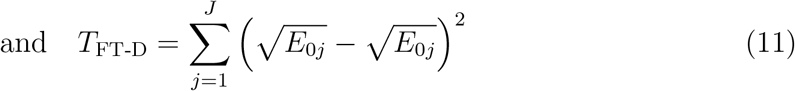

where 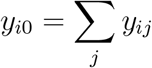 denotes the number of encounters for individual *i* across the *J* detectors and 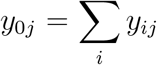 denotes the number of individuals encountered at detector *j* with 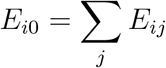 and 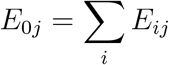.

In addition, we used Pearson’s *χ*^2^ metric (Gelman et al., 1996):

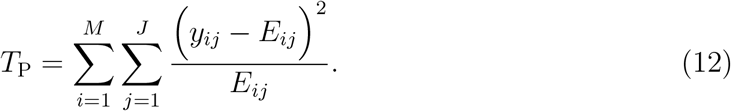

We estimated the Bayesian *p*-values for the above test quantities using a thinned sample (thinning rate = 10) of all the MCMC samples drawn during fitting of a given model.

## 3 Results

All MCMC samples of the parameters of interest (e.g., *N*, *α*% home range area, the detection function parameters) were obtained after ensuring proper mixing and convergence, with 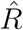 values below the nominal threshold 1.1. When fitted with the same detection function used for simulation, both population size and home range area were estimated without significant bias (average RB between 0% and 4%), with good precision (CV < 10%) and nominal levels of the coverage probability (approx. 0.95) (Fig. 2).

**Figure 2:**
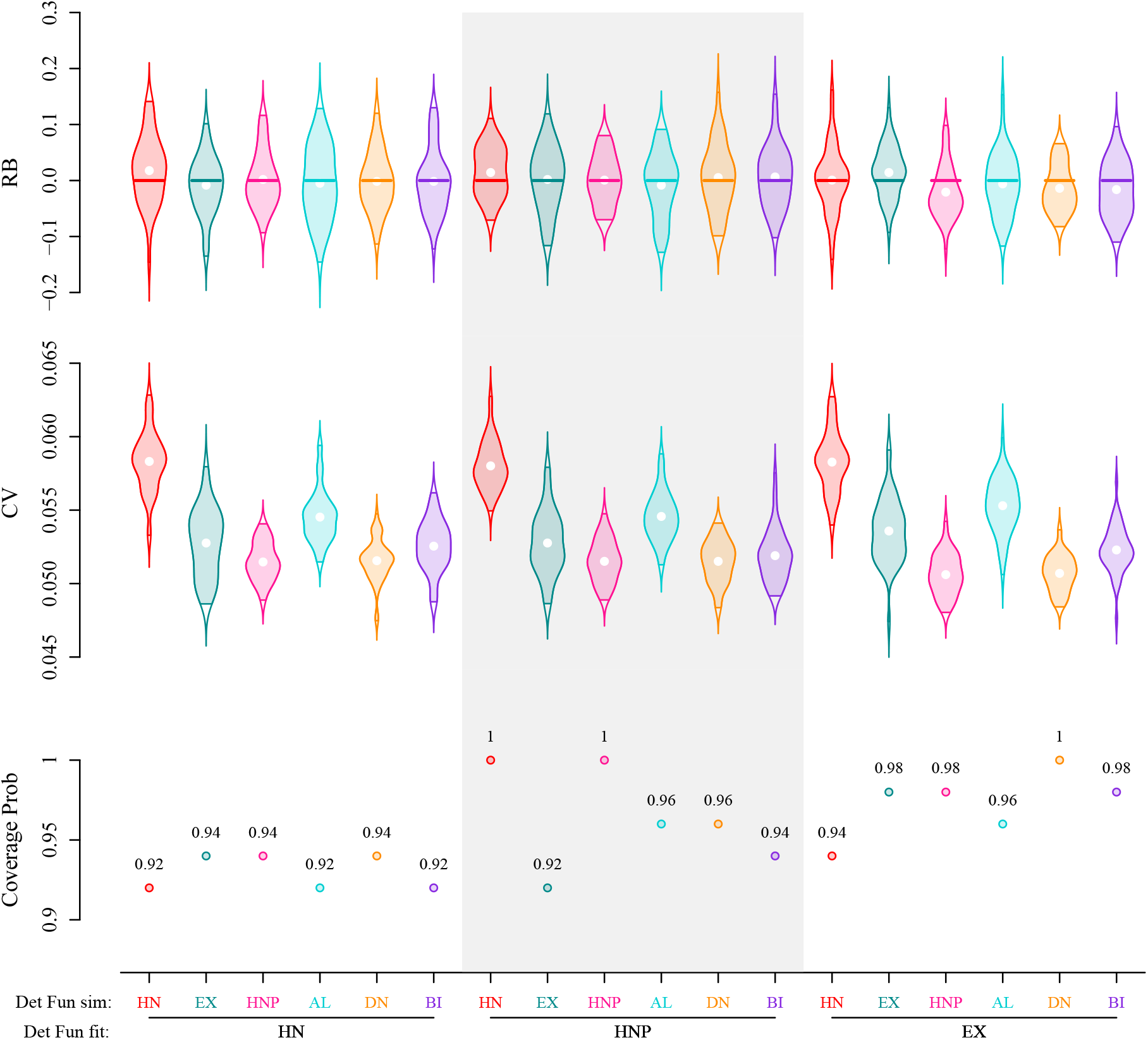
Posterior summaries of population size *N* derived using spatial capture-recapture in parameter set 1. Results compare relative bias (RB), coefficient of variation (CV), and 95% coverage probability for different pairings of simulated and fitted detection functions. Detection functions include the half-normal (HN), exponential (EX), half-normal plateau (HNP), asymmetric logistic (AL), donut (DN), bimodal (BI). Violins represent the distribution of RB / CV from 50 simulations.

### 3.1 Consequences of misspecification

#### 3.1.1 Population size

We detected no pronounced effect of the choice of detection function (HN, HNP, EX) fitted on relative bias and precision of *N*, regardless of the detection function used for simulation (Fig. 2 and Appendix Fig. 3). The average RB of *N* in parameter set 1 ranged between: −1% and 0% for model HN (avg. CV = 5.3%), between −1% and 2% for model HNP (avg. CV = 5.4%) and between −2% and 2% for model EX (avg. CV = 5.3%). Similar levels were found for parameter set 2: with average RB between −1% and 3% for model HN (avg. CV = 6.5%), between 0% and 1% for model HNP (avg. CV = 6.6%) and between −2% and 2% for model EX (avg. CV = 6.3%). Coverage probability exceeded 93% for all scenarios.

#### 3.1.2 Home range area

Misspecification of the detection function risked producing erroneous 95% kernel home range area estimates (Figs 3, 5 and Appendix Figs 4 and 6). For both parameter sets, home range area estimates were biased and had very low 95% coverage probability for the misspecified HN and EX models.

**Figure 3:**
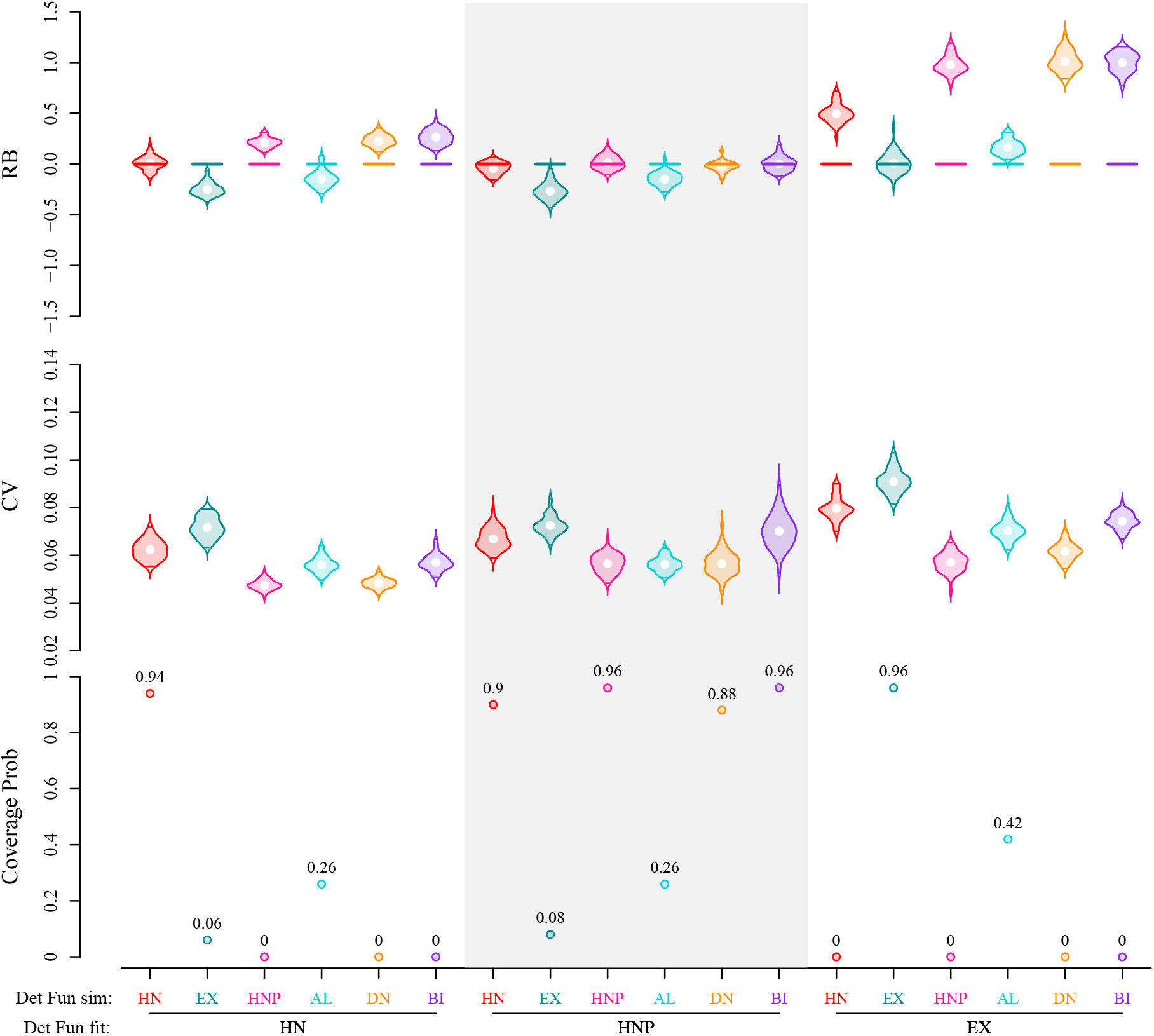
Posterior summaries of home range area derived using spatial capture-recapture in parameter set 1. Home range area was estimated as the 95% kernel of the utilization distribution from the realization of detection function used during model fitting. Results compare relative bias (RB), coefficient of variation (CV), and 95% coverage probability for different pairings of simulated and fitted detection functions. Detection functions include the half-normal (HN), exponential (EX), half-normal plateau (HNP), asymmetric logistic (AL), donut (DN), bimodal (BI). Violins represent the distribution of RB / CV from 50 simulations.

The EX model overestimated home range area by up to 170% when fitted to SCR data simulated with the HNP, DN or BI detection functions. Relative bias was also positive but generally smaller with the HN model (between 21% and 71%) when fitted to data simulated using either the HNP, DN or BI detection functions. By contrast, home range area was underestimated by the HN model when fitted to data simulated with the EX and AL detection functions for parameter set 1 (avg. RB between −14% and −25%).

The HNP model was more forgiving and accommodated data simulated with HN, DN and BI detection functions (average RB between −7% and 1%, Coverage prob *≥* 88%). Home range area estimates from data simulated with the AL detection function were better estimated for parameter set 2 (average RB 7%, coverage prob. = 78%) than for parameter set 1 (average RB −15%, coverage prob. = 26%). However, HR estimates were highly biased (average RB between −31% and −26%) with low coverage probability (*≤* 10%) when fitted to the data set simulated with the EX model.

### 3.2 Goodness of fit

Bayesian *p*-values based on individual and detector level counts (*T*_FT-I_ and *T*_FT-D_) were all centered on 0.5 but highly dispersed, thus failing to reveal any misspecification in detection functions. On the other hand, Bayesian *p*-values targeting discrepancy in observations at both the individual and detector level, corresponding to Pearson’s *χ*^2^ and Freeman-Tukey’s (FT) metrics were more useful for detecting a lack of fit (Fig. 4 and Appendix Fig. 5). First, and as expected, they showed high concentration around 0.5 (no evidence of lack of fit), when the fitted detection function matched the one used for simulations. Taking both parameter sets into account, Bayesian *p*-values *p*(*χ*^2^) were estimated with means 0.49 (HN model, CV 19.9%), 0.51 (HNP model, CV 12.2%), 0.50 (EX model, CV 26%); and *p*(FT) were estimated with means 0.49 (HN model, CV 5.1%), 0.49 (HNP model, CV 3.4%), 0.50 (EX model, CV 4.4%).

**Figure 4:**
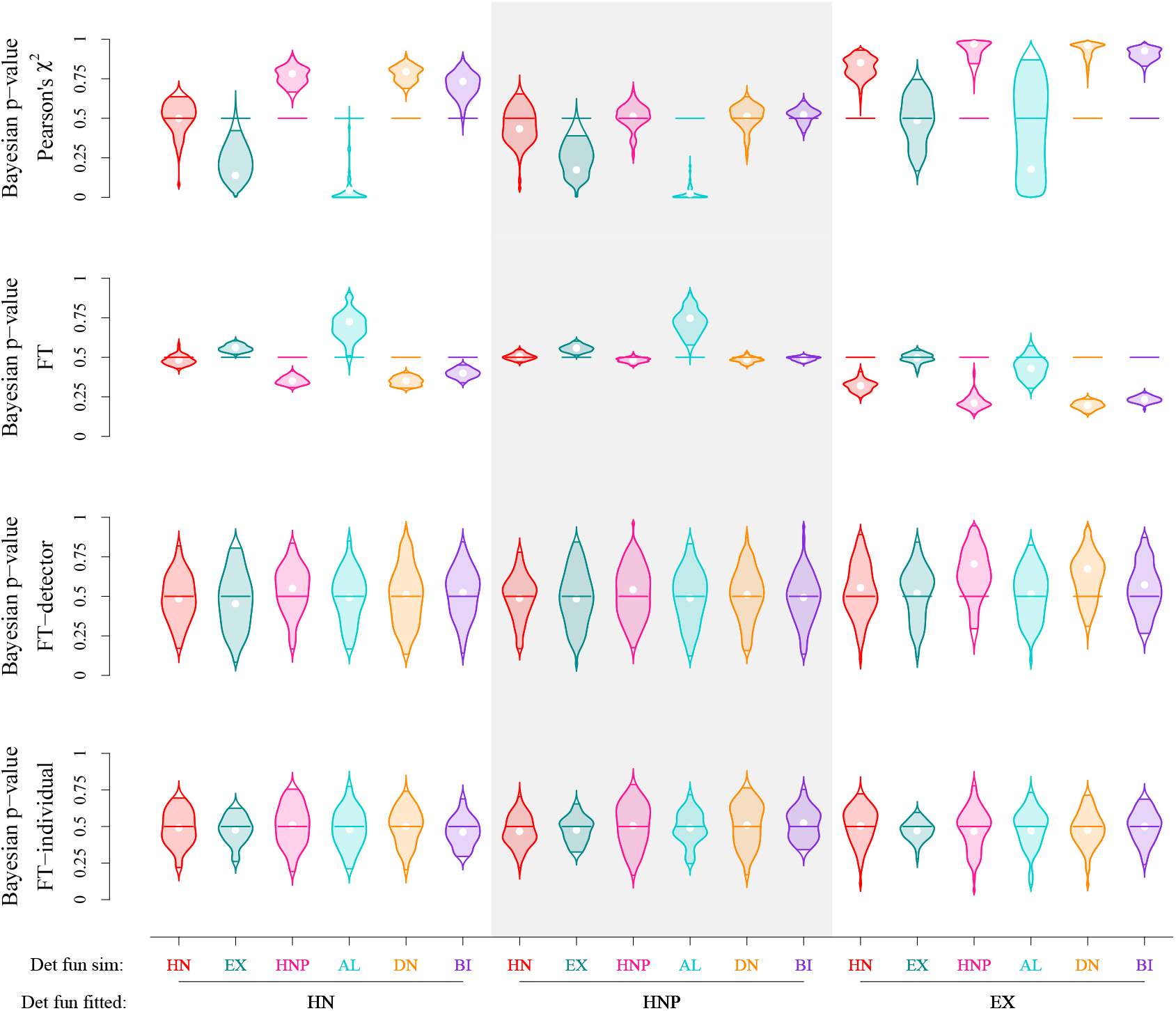
Estimates of Bayesian *p*-values from different metrics: Freeman-Tukey (FT), Pearson’s *χ*^2^, FT metric based on individual level count (FT-I) and FT metric based on detection level count (FT-D). Graphs compare the Bayesian *p*-value estimates between different pairings of simulated and fitted detection functions for parameter set 1. Each Violin represents the distribution of Bayesian *p*-values for a specified metric from 50 simulations.

**Figure 5:**
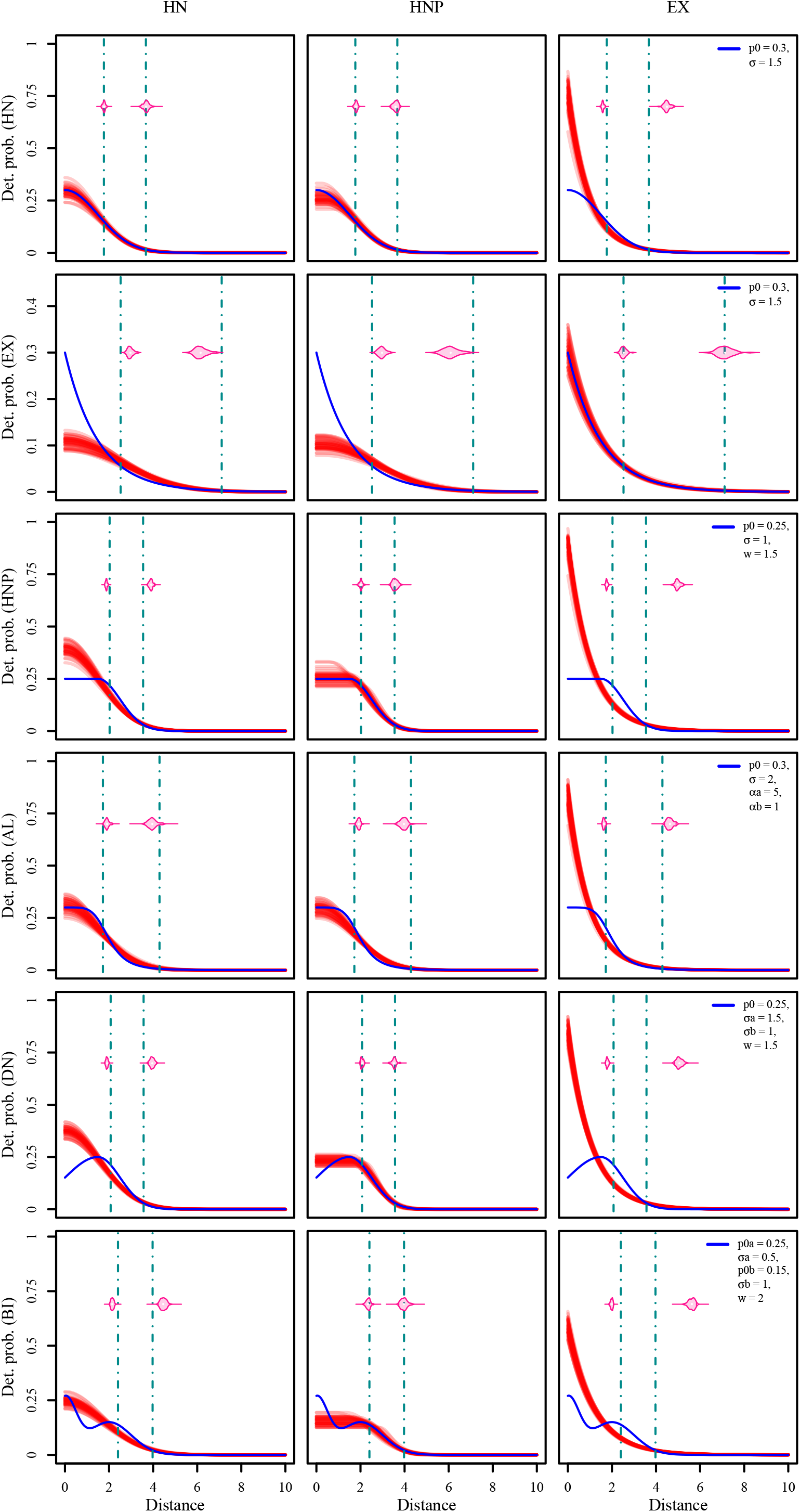
Comparison of estimated detection functions (‘red’ lines) and the estimates of home range radius (50% and 95% quantiles, ‘pink’ violins representing the distribution from 50 simulations) with the ‘true’ detection function (‘blue’ line) and ‘true’ home range radius (‘deep cyan’ vertical lines) for different scenarios in parameter set 1. Rows correspond to the ‘true’ detection function that was used to
fit the SCR model. Parameter estimates of each model fitting were used to estimate the fitted detection function and they are plotted as a function of distance in arbitrary distance units.

Further, these metrics showed the greatest departure from 0.5 when the detection functions were misspecified for parameter set 1. Whereas, for parameter set 2 (larger home range areas and overlap), the detectability of a lack of fit was comparatively poor in case of the misspecification (95% CI for *p*(*χ*^2^) = (0.26, 0.90), for *p*(FT) = (0.25, 0.53)). The most pronounced indication of lack of fit was observed when fitting the exponential model to data simulated with detection functions other than exponential (95% CI for *p*(*χ*^2^) = (0.01, 0.99), for *p*(FT) = (0.16, 0.50) in parameter set 1; 95% CI for *p*(*χ*^2^) = (0.63, 0.92), for *p*(FT) = (0.22, 0.49)) for parameter set 2).

Bayesian *p*-values (Pearson’s *χ*^2^, FT) clustered around 0.5 for the HNP model fit to data simulated with the HN, DN and BI detection functions. Bayesian *p*-value estimates were near 0 for both the EX or AL model fit to data simulated with the HN, DN and BI detection functions, indicating a pronounced lack of fit. SCR data sets simulated using the EX and AL detection functions, due their longer right tails, occasionally resulted in very distant detections (*>* 9 du) from individual ACs. SCR models with HNP and HN detection models have difficulties to accomodate these distant detections. This is reflected in the posterior samples of detection probability for these detections, being of infinitesimal magnitude under HNP model and hence minuscule Bayesian *p*-value estimates *p*(*χ*^2^).

## 4 Discussion

Our study revealed that misspecification of the detection function in SCR can have consequences for inference that range from benign to severe. While population size estimates are robust, inference about space use can be highly impacted by the wrong choice of detection function. Fortunately, extreme misspecification, with the strongest impact on parameter estimates, are also the ones most readily discovered using Bayesian *p*-values. We also found that some detection functions are better able to accommodate a variety of space use patterns than others, with the half-normal plateau and exponential function being the most and the least flexible, respectively.

### 4.1 Consequences

We tested the consequences of misspecifying the detection function for a wide range home range utilization distributions (Fig. 1). We found little effect of misspecifications on the precision and accuracy of population size estimates. This corroborates findings from other studies that population size estimates from SCR models are generally robust to misspecification of the detection function (Efford, 2004). Recently, Efford (2019) showed with simulations that SCR estimates of population size were also robust to non-circularity of home ranges (e.g., elliptical, rectangular) as long as recaptures were allowed in both directions of the detector array.

Contrastingly, all misspecifications that we tested led to pronounced bias in home range area estimates when fitted with the exponential or half-normal models. Relative bias was especially severe when using a misspecified exponential model (*>* 100% in parameter set 2) and, to a lesser degree a half-normal model (*≈* 35% in parameter set 2). This finding is of concern, as the half-normal is by far the most commonly used detection function in SCR analyses (Royle et al., 2014). It is also noteworthy that a simple expansion of the half-normal, the half-normal plateau detection function proved to be the most accommodating when fitted to data with contrasting space use patterns. This is presumably due to the extra parameter (plateau width) which provides substantial additional flexibility. Only when fitted to data simulated using the AL and EX did the HNP led to significant bias in home range area estimates.

### 4.2 Diagnostics

We found that misspecifications of the detection function can be challenging to detect using standard goodness of fit diagnostic Bayesian *p*-values. Among the four metrics used, the FT and Pearson’s *χ*^2^ were the only ones which were able to reveal missspecifications in the detection function (Section 3.2). The general recommendation to identify a lack of fit in SCR is based on whether or not the Bayesian *p*-value falls outside the interval (0.1, 0.9) (Royle et al., 2014). Using this threshold, the Bayesian *p*-values would have often failed to reveal the missspecification. Indeed a *p*-value of 0.25 or 0.75 does not guarantee that the detection function is correctly specified and that home range area estimates are free from bias. We found a positive correlation between the RB and *p*(*χ*^2^) (Appendix Fig. 7), meaning that a model with a Bayesian *p*-value *>* 0.5 is likely to overestimate home range area, while a model with a Bayesian *p*-value < 0.5 is likely to underestimate it (Appendix Fig. 7). For example, HN models fitted to SCR data simulated with the HNP detection function led to an average *p*(*χ*^2^) of 0.78 (with 2.5% and 97.5% quantiles as 0.67, 0.88 respectively) and to overestimated home range area estimates (21% average relative bias with 0% coverage prob., Fig. 4). Conversely, HN models fitted to SCR data simulated with the AL detection function led to an average *p*(*χ*^2^) of 0.04 (with 2.5% and 97.5% quantiles as 0, 0.31 respectively, see Fig. 4) and to underestimated home range area estimates (−14% average relative bias with 26% coverage prob.). However, a *p*(*χ*^2^) value of 0.08 could also be obtained from a correctly specified HN model. This illustrates the ambiguity associated with the use of a single Bayesian *p*-value as an indicator of model misspecification and erroneous home range area estimates (Stern and Cressie, 2000). The use of *p*-value has been criticized in the literature for being conservative due to apparent ‘double use’ of the data (Dey et al., 1998; Bayarri and Berger, 2000).

Despite the relatively low power of the GOF metrics considered here to detect a lack of fit, it is worth noting that the most problematic deviations could still be identified. For instance, in extreme cases of model misspecification (e.g., when using a misspecified SCR model with an EX detection function) where home range space use considerably differed from simulated ‘true’ space use pattern (leading to highest relative bias and lowest coverage values), Bayesian *p*-values were furthest from 0.5 and were thus effective in identifying the lack of fit (Appendix Figs 7 and 8). We argue that rejection of models with moderate magnitude of Bayesian *p*-values should depend on the aim of the study (parameters targeted) rather than blindly following the rule of thumb (Stern and Cressie, 2000).

Practitioners should choose the Bayesian *p*-value discrepancy metric with caution. As we have shown, different metrics perform differently. The FT metric with variance stabilising property was overly optimistic in assessing the goodness of fit and showed high concentration around 0.5, even in cases of severe model misspecification. The pooled version of the discrepancy metrics (e.g., FT-I and FT-D) showed extremely high variance (Fig. 4) which may be similarly misleading about the quality of model fit. Appropriate goodness of fit metrics should be chosen based on their ability to detect certain misspecifications, instead of selecting and reporting those metrics that indicate a good fit (Head et al., 2015).

### 4.3 Recommendations

Based on the results from the simulation study, we can make recommendations for SCR users regarding the choice of a detection function. Here it is worth repeating that density and population size estimates are largely immune to misspecifications in the detection function. However, if the goal is to estimate animal space-use, SCR users need to be more cautious.

We recommend fitting multiple SCR models with different detection functions and comparing their estimates and associated Bayesian *p*-values. Even though the relative bias and coverage values cannot be assessed when fitting models to empirical data, comparing estimates and *p*-values among models can be informative, i.e. by revealing which of the candidate models most closely approximates the space use patterns in the study system.

Our results suggest that the highest risk of flawed inferences is associated with the exponential detection function. Consider two hypothetical scenarios. First, one can fit all three detection functions tested here to an SCR data set where the true, but unknown, detection function is the exponential one. In this situation, both the mean estimates for home range area and the Bayesian *p*-values for the SCR models fitted with HN or HNP detection function are likely to be lower than the ones from the model fitted with the exponential function (Fig. 4). This would identify the exponential function as the most likely candidate detection models (among those considered) and thus avoid under-estimating home range area. Second, one can fit all three models to another SCR data set where the true, but unknown, detection function is not the exponential one. This will likely lead to larger Bayesian home range area estimates and *p*-values with the exponential model compared to the two others functions, thus ruling out the exponential function as the data-generating process and avoiding the associated large positive bias (Fig. 3). If in doubt, we recommend using the half-normal plateau detection function, as it is flexible enough to accommodate most home range shapes tested here and return sensible home range area estimates. However, fitting the half-normal plateau detection function comes at the cost of estimating an additional parameter *w* and has proved to be more challenging to fit than the simpler half-normal or exponential functions. Finally, alternative sources of information, such as telemetry data, may allow independent estimation of individual space-use pattern in the study population (Aarts et al., 2008; Kie et al., 2010; Royle et al., 2013b) and inform the choice of detection function.

Concerning GOF, neither detector nor individual-level metrics were able to identify misspecifications of the detection function in our study. We instead recommend the use of discrepancy metrics computed at the level of the individual *and* detector-specific detections in order to help reveal mismatch between the detection function and individual space use.

### 4.4 Conclusions

Although initially developed with the intent to provide spatially referenced estimates of population size (Efford, 2004; Efford et al., 2004; Borchers and Efford, 2008), SCR is increasingly being used for estimating parameters that inform about other ecological aspects (Chandler and Clark, 2014; Bischof et al., 2016; Muneza et al., 2017; Ergon and Gardner, 2014; Chandler et al., 2018). SCR holds particular promise for addressing spatial ecological questions at the population level (Morin et al., 2017; Bischof et al., 2017). Whereas studies focusing exclusively on population size and density estimates may only be marginally impacted by misspecified detection functions (our results and those by Efford and Dawson, 2009; Russell et al., 2012), the repercussions can be severe when inferences about space use, e.g. home range area, are sought. It is in these kinds of studies where we expect the greatest benefit from combining simulations and GOF for selecting a suitable detection function. Here, we used the GOF statistics to identify missspefications in the shape of the detection functions, however, missspecification of spatial and individual covariates on detection probability is also likely to occur in monitoring studies. We emphasize that assessing the goodness of fit is an integral part of data analysis and urge for developing custom model checking diagnostics for testing different source of misspecification in Bayesian SCR models.

## Acknowledgements

This work was funded by the Norwegian Environment Agency (Miljødirektoratet), the Swedish Environmental Protection Agency (Naturvårdsverket), and the Research Council of Norway (NFR 286886). Computation was performed on resources provided by Uninett Sigma2 - the National Infrastructure for High Performance Computing and Data Storage in Norway.

## Authors contribution

S.D and R.B developed the concept and methodology. S.D led the analysis with help from C.M, P.D and R.B to run the models. All authors S.D, C.M, P.D and R.B contributed writing of the manuscript and gave final approval for publication.

## Data accessibility

No empirical data were used in this analysis. R code for generating simulated data and a few additional figures are provided in the Supplementary material Appendix.

